# SARS-CoV-2 infection activates inflammatory macrophages in vascular immune organoids

**DOI:** 10.1101/2024.03.20.585837

**Authors:** Chiu Wang Chau, Alex To, Rex K.H. Au-Yeung, Kaiming Tang, Yang Xiang, Degong Ruan, Lanlan Zhang, Hera Wong, Shihui Zhang, Man Ting Au, Seok Chung, Euijeong Song, Dong-Hee Choi, Pentao Liu, Shuofeng Yuan, Chunyi Wen, Ryohichi Sugimura

## Abstract

SARS-CoV-2 provokes devastating tissue damage by cytokine release syndrome and leads to multi-organ failure. Modeling the process of immune cell activation and subsequent tissue damage is a significant task. Organoids from human tissues advanced our understanding of SARS-CoV-2 infection mechanisms though, they are missing crucial components: immune cells and endothelial cells. This study aims to generate organoids with these components. We established vascular immune organoids from human pluripotent stem cells and examined the effect of SARS-CoV-2 infection. We demonstrated that infections activated inflammatory macrophages. Notably, the upregulation of interferon signaling supports macrophages’ role in cytokine release syndrome. We propose vascular immune organoids are a useful platform to model and discover factors that ameliorate SARS-CoV-2-mediated cytokine release syndrome.

## Introduction

COVID-19 pandemic of the last few years challenged health, social, and humane communities. Lethality with the SARS-CoV-2 infection is marked by multi-organ failure by cytokine release syndrome and resultant vascular damage ^1–3^. Since the beginning of the pandemic, major focuses were put in animal models, patient samples, and autopsies to identify the key mechanisms involved in ^4–6^. Among the experimental models, organoids gained great attention due to its capacity to recapitulate tissue structure and complex cell-cell interactions, multiple organoids identified receptors, infection mechanisms, and propagation of virus^7–13^. However, as organoids are made largely from epithelial cells of tissues, macrophages and vasculatures are usually missing^14^. Because macrophages are a major mediator of cytokine release syndrome and damage vasculature, the lack of these effectors and targets hampers key knowledge from organoid models^15–17^. Having vascular networks and immune cells in organoids is a promising venue^18^. We previously demonstrated the generation of vascular immune organoids (VIOs) to model macrophage developments^19^. Here we generated organoids composed of vascular networks and macrophages and examined the response to SARS-CoV-2 exposure. We detected expression of SARS-CoV-2 receptors, inflammatory macrophage activation, and potential indicators of cytokine release syndrome. This platform allows the assessment of how macrophages produce cytokine storm to cause vascular damages. Such vascular immune organoids will help us understand mechanisms and discovery of therapeutics to prevent lethality from the infection.

## Results

To examine the effect of infection of SARS-CoV-2 in the vasculature and mesenchymal tissues, we generated vascular organoids from human pluripotent stem cells. We differentiated endothelial progenitor cells via Stemdiff Hematopoioetic kit and mesenchymal stromal cells via Ascorbic acid and Activin A respectively. We then aggregated them to form a 3D structure (**Figure 1A**). We named the resultant structure the vascular organoids (VO) featured by the balloon-like transparent area covered with dark streaks indicative of vessels **(Figure 1B**). We detected the vascular network by both paraffin section and whole-mount staining with endothelial marker CD31 (**Figures 1C-D**). To determine SARS-CoV-2 infection and its effect, we infected VO (**Figure 1E**). We observed cell death on the surface of VO indicating the virus’ toxic effect (**Figure 1F**). We observed an increased cleaved caspase-3 signals in infected VO, indicating apoptosis **(Supplementary figure 1A-C)**. We also detected the Von Willebrand factor (vWF) after infection, indicating vascular activation after injury **(Supplementary figure 2A)**. We then detected viral N-protein in infected VO, indicating successful infection (**Figure 1G**). These data demonstrate that VO can be infected with SARS-CoV-2 and provokes cell death.

**Figure 1.**
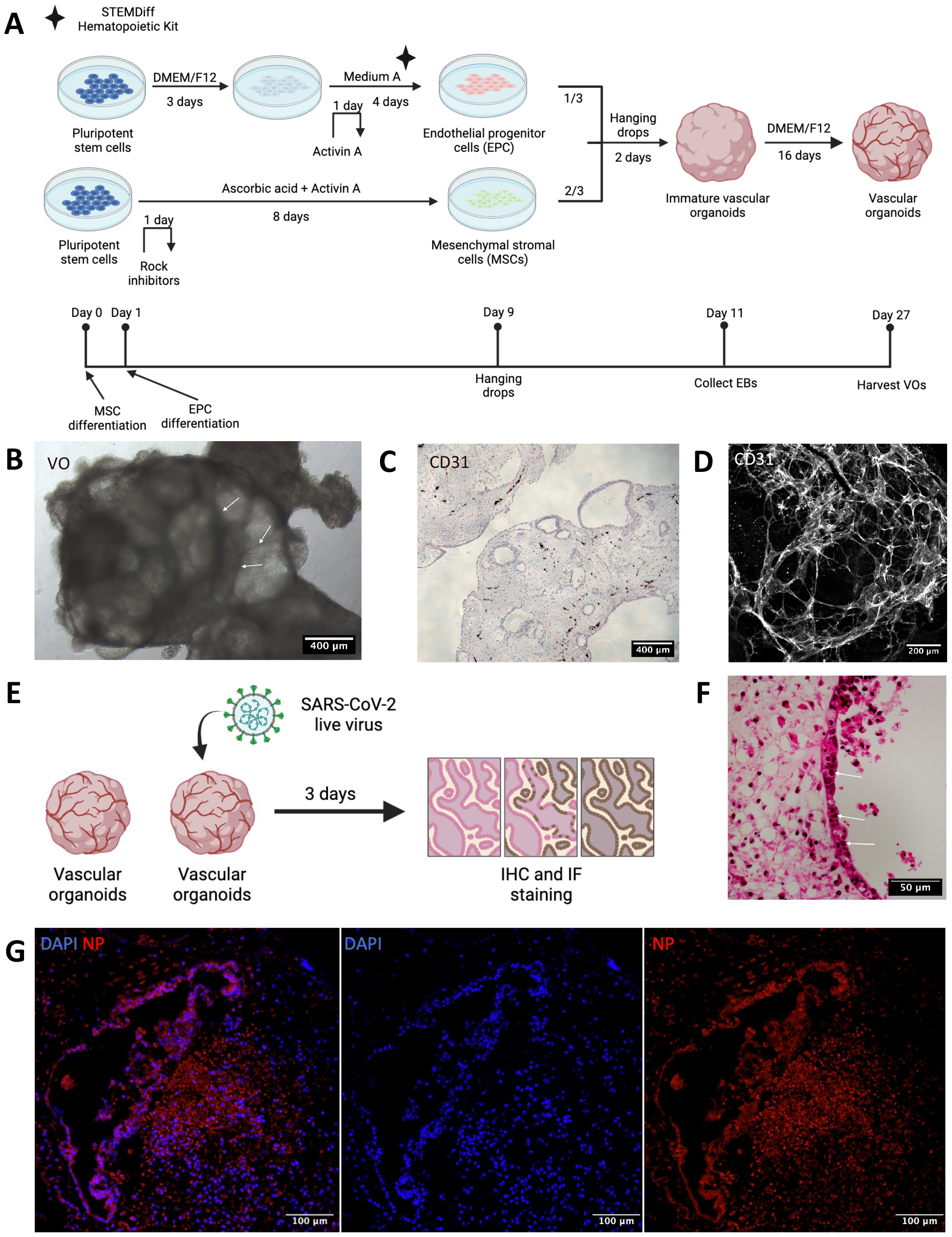
Vascular organoids model SARS-CoV-2 infections. **(A)** The scheme of vascular organoids (VOs) generation. hPSCs are differentiated into endothelial progenitor cells (EPCs) by STEMdiff hematopoietic kit medium A (i.e. supplement A), and into mesenchymal stromal cells (MSCs) by ascorbic acids and activin A independently. Mature MSCs and EPCs are eventually aggregated to generate vascular organoids (VOs). **(B)** Brightfield microscopy image of VOs. Examples of dark streaks indicative of vessels are highlighted by arrows. Scale bar = 400 μm. **(C)** CD31 IHC staining of VO. Scale bar = 400 μm. **(D)** CD31 whole-mount immunofluorescent staining of VO. Scale bar = 200μm. **(E)** The scheme of SARS-CoV-2 infection to VO. Briefly, 1E7 PFU/mL of SARS-CoV-2 live virus was loaded to infect VOs for 72 hours. **(F)** H&E staining indicates cell death over the surface of VO after infection. Examples of nuclear inclusion bodies are highlighted by arrows. Scale bar = 50 μm. **(G)** Immunofluorescent staining shows a successful infection by the presence of viral N-protein (NP) in infected VO. Scale bar = 100 μm.

**Figure 2.**
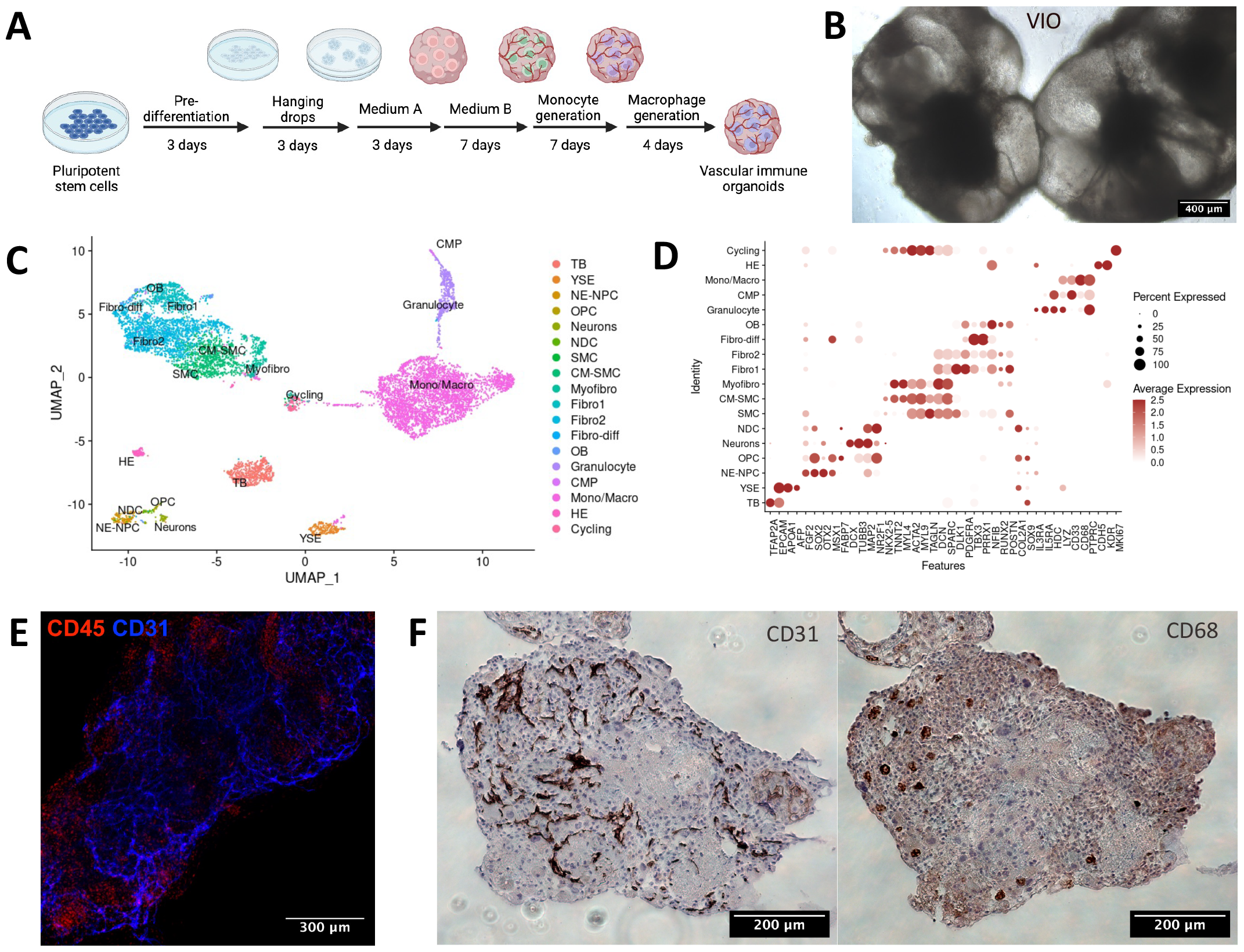
Vascular immune organoids co-develop vasculature and macrophages. **(A)** The scheme of vascular immune organoid (VIO) generation. **(B)** Bright-field microscopy image of VIO. Scale bar = 400μm. **(C)** UMAP clustering of VIO from scRNA-seq shows vascular endothelium, myeloid cells, and fibroblasts. **(D)** Dotpots of representative markers in each UMAP clusters. **(E)** Whole-mount immunofluorescent staining of CD31 (blue) and CD45 (red) of VIO. Scale bar = 300 μm. **(F)** CD31 and CD68 IHC staining of paraffin, uninfected VIO. Scale bar = 200μm. (Left: CD31; Right: CD68).

To further examine SARS-CoV-2 mediated inflammation by immune cells, we modified VO to co-develop immune cells. We added hematopoietic cytokines during vascular organoid formation (**Figure 2A**). The resultant organoid generated hematopoietic cells and is called vascular immune organoids (VIO) hereafter. VIO showed a transparent balloon-like structure like VO, indicative of mesodermal cells emergence and migration (**Figure 2B**). VIO comprises vascular endothelium, myeloid cells, and fibroblasts as shown by single-cell RNA sequencing (**Figure 2C, 2D**). The whole-mount immunofluorescence detected a cluster of CD45+ hematopoietic cell areas vascularized by CD31+ endothelial cells (**Figure 2E**). CD31+ endothelial network and CD68+ macrophages are distributed throughout VIO (**Figure 2F**). These data demonstrated that VIO possessed macrophages and vasculature, which could be applied to the SARS-CoV-2 infection model.

We then verified the expression of Toll-like receptor (TLR) 4, a known receptor of SARS-CoV-2^20–23^. We detected TLR4 in both monocytes and endothelial cells at the messenger level by single-cell RNA-sequencing analysis and protein level by flow cytometry (**Figures 3A-B**). We then examine the expression of other known SARS-CoV-2 receptors^24–28^. Other receptors such as ACE2 are absent in both endothelial cells and monocytes and rather expressed in miscellaneous trophoblast-like cells. The absence of ACE2 in endothelial cells are consistent with other papers^20,29,30^. BSG^31^ and NRP1^26,32^ expressed endothelial cells, stromal cells, and monocytes. TMPRSS2^27^ was nearly absent in the organoids (**Figure 3C**). We further detected NP in VIO, showing successful SARS-CoV-2 infection **(Supplementary figure 2B)**. These data indicate that VIO expresses receptors of SARS-CoV-2 and are successfully infected.

**Figure 3.**
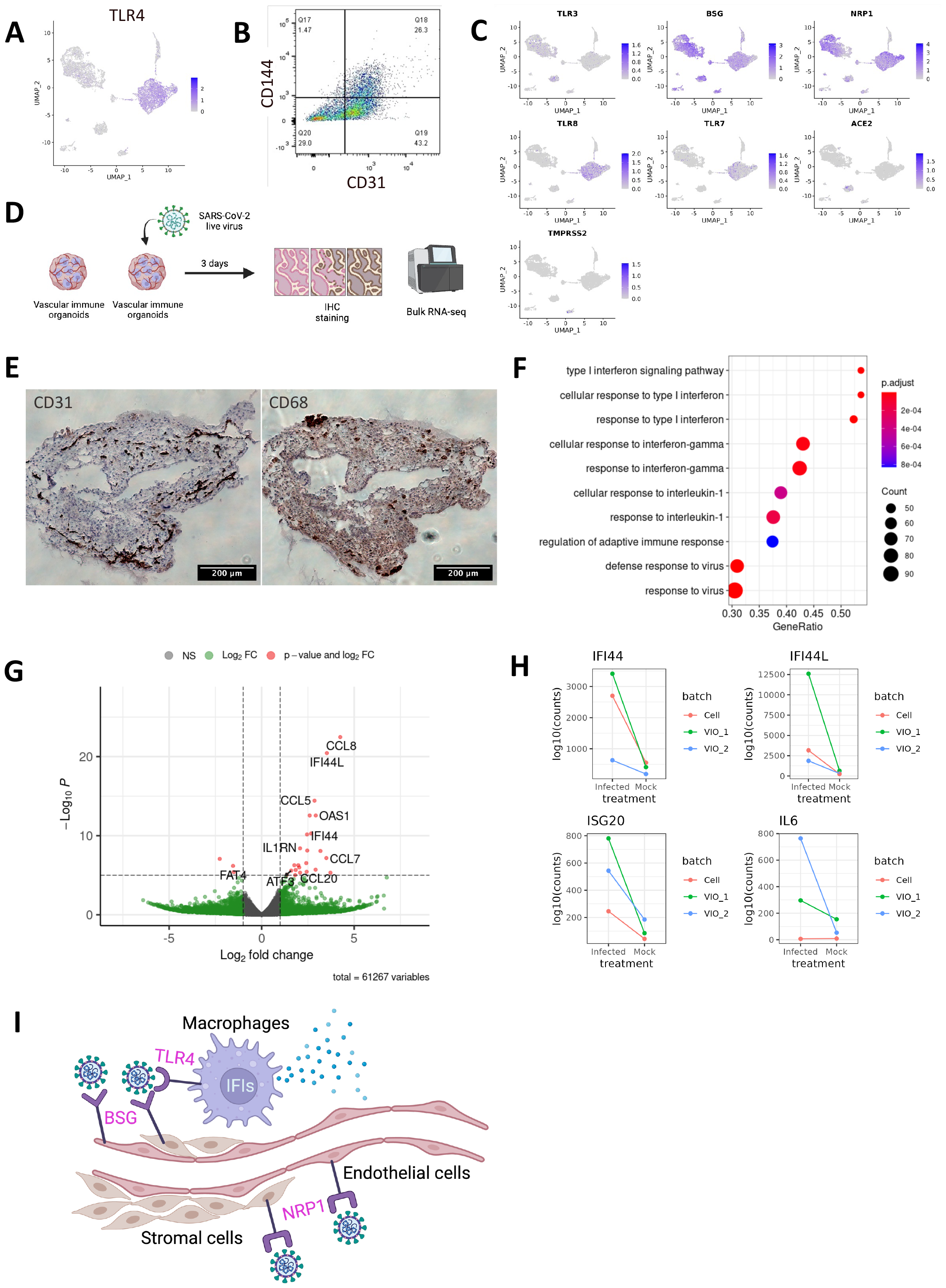
SARS-CoV-2 infection activates inflammatory responses. **(A)** Single-cell RNA expression of TLR4 in VIO. **(B)** Surface TLR4 (CD133) expression by flow cytometry of VIO. **(C)** Single-cell RNA expression of viral receptors in VIO. **(D)** The scheme of SARS-CoV-2 infection to VIO and subsequent bulk-RNAseq experiments. **(E)** CD31(left) and CD68(right) IHC staining of infected VIO. Scale bar = 200 μm. **(F)** Gene set enrichment analysis (GSEA) result of infection vs mock VIOs. **(G)** Volcano plot of differential expressed genes of infected vs mock VIOs. **(H)** RNA expression of IFI44, IFI44L, ISG20, and IL6 in infected versus mock VIO and isolated immune cells. **(I)** A proposed model of SARS-CoV-2 pathogenesis in VIO.

To determine if virus infection provoked inflammatory programs, we did bulk RNA-sequencing of VIO infected with SARS-CoV-2 compared with uninfected ones (**Figure 3D**). After virus infection, CD31+ endothelium and CD68+ macrophages remained. Notably, CD68+ macrophages co-localized on the surface of VIO, suggesting an inflammatory reaction by virus exposure (**Figure 3E)**. We identified that infected VIO upregulated inflammatory signatures. These are primarily transduction and response to Type I Interferon, Interferon Gamma, and response to viruses (**Figure 3F)**. We further verified macrophage-mediated inflammation by upregulation of genes IFI family (interferon-induced inflammation), and CCL family (macrophage chemotaxis) (**Figure 3G**). We next detected a pro-inflammatory cytokine IL6 in infected VIO **(Supplementary figure 2C)**. This demonstrates VIO can potentially be used to study cytokine release syndrome.

To show macrophage-specific response to SARS-CoV-2, we collected released monocytes from VIO and differentiate them into macrophages. We next infected the isolated macrophages by SARS-CoV-2 and performed bulk RNA-seq **(Supplementary figure 3A)**. We then integrated the VIO and isolated immune cells data for downstream bulk RNAseq analysis. We observed that IFI family and CCL family are consistently upregulated **(Supplementary figure 3B)**. We observed mostly interferon-related responses, NF-kB, TNF, and JAK-STAT pathways after SARS-CoV-2 infection **(supplementary figure 3C-D)**. The integrated analysis demonstrate mostly macrophage-specific response to SARS-CoV-2 **(Supplementary figure 4)**. We finally showed infected VIO and isolated macrophage consistently upregulated IFI44, IFI44L, and ISG20 expression **(Figure 3H)**, which are known as the interferon responses to defence against virus. However, a pro-inflammatory gene like IL6 is regulated only in infected VIO but not in isolated macrophages **(Figure 3H)**. This indicate the important roles of other cell types contributing to the cytokine release syndrome, and justify the usage of VIOs to study the SARS-CoV-2 pathogenesis. These data demonstrate that SARS-CoV-2 infection provoked macrophage inflammation. Macrophages exposed to SARS-CoV-2 activated innate immune programs including interferon expression and upregulated downstream pathways. Taken together, VIO could reveal the role of macrophages and other cell types in inflammatory reaction to SARS-CoV-2 infection **(Figure 3I)**.

## Discussion

In this study, we established vascular immune organoids (VIOs) and demonstrated that infection with SARS-CoV-2 activated inflammatory macrophages as showed by IFI family (interferon-induced inflammation) and CXCL family (macrophage chemotaxis) (figure 3G). Unlike other vascular organoids from human pluripotent stem cells^33,34^, our VIOs possess hematopoietic cells including macrophages which is able to induce interferon induced response an anti-viral defence response (GSEA) and produce IFI and ISG genes (figure 3F, H). This suggests that VIOs serve as a platform for understanding the mechanisms of SARS-CoV-2-mediated cytokine release syndrome. As such syndrome and subsequent vascular damage lead to multi-organ failure in COVID-19 lethality, we propose that VIO could be suitable for therapeutic discovery to ameliorate the syndrome. The current study is at this point limited to vasculature and macrophages. Including lymphoid lineage by cytokines presumably IL-7 and IL-15 with longer-term incubation would recapitulate the infection and downstream inflammatory events more precisely. The ultimate application of VIO would be to integrate with other tissue organoids and serve both vascular networks and various immune cells. This assembloid strategy would provide a significant venue not only for SARS-CoV-2 but also for other infections and pathogen modelings.

## Data availability

The bulk RNA-seq data of mock and infected vascular organoids are deposited in the Sequence Read Archive (SRA) under accession number PRJNA1051064. The proceeded expression data is available in GitHub (https://github.com/alexto22/COVID19_VIO). The scRNAseq data of vascular immune organoids is deposited in Gene Expression Omnibus (GEO) under accession number GSM6755678 (https://www.ncbi.nlm.nih.gov/geo/query/acc.cgi?acc=GSM6755678).

## Author contribution

C.W.C. and A.T. designed, conducted experiments and analyzed data. R.K.H.A.Y. performed IHC staining and analysis. K.T., Y.S., and C.W. conducted and supervised virus infection. Y.X., D.R., L.Z., H.W., M.T.A. helped organoid formation. S.Z. guided scRNA-seq analysis. S.C., E.S., and D.H.C. assisted image analysis. P.L., C.W., and R.S. conceived the projects. C.W.C., A.T., and R.S. wrote the manuscript. All the authors discussed the manuscript.

## Acknowledgements

We would like to thank CPOS HKU for the assistance in flow cytometry, tissue sectioning, and next-generation sequencing. R.S. is supported by Seed Funding from HKU, RGC ECS 27109921, RGC GRF 17109022, and the InnoHK Centre for Translational Stem Cell Biology.

## Methods

### Human pluripotent stem cell culture

Human C5-expanded potential stem cells was developed by Pentao Liu’s lab^35^ and was used as human pluripotent stem cells (hPSCs) in this paper. hPSCs are cultured in a feeder-based system. SNL feeders are seeded onto gelatine-coated 12-well plates 2 days before hPSC passage. hPSCs are cultured with maintenance medium as described^35^. Medium is changed every day until 70% confluence. hPSCs are dissociated with 30 mM EDTA/ PBS and split at 1:6 – 1:8 ratio.

### Vascular organoids (VOs) formation

#### Mesenchymal stromal cells differentiation

hPSCs are dissociated with 30 mM EDTA/PBS on day 1 and seeded onto gelatin-coated 6 well plates at 60k cells/well in 1 mL mesenchymal differentiation medium (10% KOSR/DMEM/F12/100 uM ascorbic acid/Activin A) supplemented with 10μM rock inhibitor. 1 mL of mesenchymal differentiation medium is added after 24 hours and medium is half changed on day 3,6,9.

#### Endothelial progenitor cell differentiation

70% confluent hPSCs are medium changed on day 0 and cultured with 10% KOSR/DMEM/F12 for pre-differentiation for 3 days. On day 3 the medium is replaced with Stemdiff hematopoietic kit medium A supplemented with 1ng/ml activin A. Medium is changed on day 4 to medium A without activin A

#### Organoid aggregation and differentiation

Mesenchymal stromal cells (MSCs) were dissociated by TrypLE while endothelial progenitor cells (MPCs) are dissociated with 30 mM EDTA/PBS. Cell clusters were gently pipetted to dissociate cells into single cells. Using hemocytometer to count cells, MSCs and EPCs are mixed in 2:1 ratio and concentration is adjusted to 20k cells / 25 uL in MSC differentiation medium supplemented with 10μM Rock inhibitor. Hanging drops are made using 25 uL cell suspension each and collected after 2 days into a 24-well low adherent plate and cultured with organoid culture medium (10% KOSR/DMEM/ 100 uM ascorbic acid).

### Vascular immune organoids (VIOs) formation

We generated VIO according to our previous studies^36,19^. In brief, we used Stemdiff Hematopoietic kit for hematoendothelial specification, followed by Stemdiff Monocyte kit for monocyte differentiation, and M-CSF for macrophage differentiation.

### Wholemount immunofluorescence

Organoids are first rinsed with PBS and fixed with 4% PFA at 4°C for 30 minutes. Then the organoids were permeabilized with staining buffer (5% FBS/ 0.5% Triton X 100/ PBS) overnight at 4°C. Primary antibodies of CD31 (Abcam, ab182981) and CD45 (Abcam, ab8216) were used at 1:200 dilution in staining buffer and incubated in dark overnight at 4°C. Secondary antibodies of and anti-mouse (Abcam, AB175473) and anti-rabbit (Abcam, AB150077) were stained at 1:400 dilution with 0.5 uM DAPI at 4°C overnight after washing 3 times with staining buffer for 15 minutes each. Organoids are washed 3 times in staining buffer for 15 minutes each and immersed in 70% glycerol/PBS for 24h before imaging. Confocal immunofluorescence images were taken by Zeiss LSM 800 or Zeiss LSM 900 at HKU CPOS.

### Paraffin sections immunofluorescence

Fixed organoids are embedded in paraffin and sectioned by HKU histopathology service by Department of Pathology. Paraffin sections are dewaxed and rehydrated by going through a ladder of xylene, ethanol and distilled water as described. Sections are then blocked and permeabilized with section staining buffer (5% FBS/0.2% gelatin/0.25% Triton X 100/PBS) for 30 minutes. Primary antibodies of anti-SARS/SARS-CoV-2 Nucleocapsid Monoclonal Antibody (Invitrogen, MA1-7404) is diluted in section staining buffer at 1:400 and sections are stained at 4°C overnight. Sections are then washed with section staining buffer 3 times for 15 minutes each. Sections are then stained with secondary antibodies (Abcam, AB175473) diluted at 1:800 and 0.25 uM DAPI at 4°C overnight. After 3 rounds of washing, direct conjugated antibodies are stained at 1:400 dilution at 4°C overnight. Slides are washed 3 times with staining buffer before mounting in 70% glycerol/PBS and sealed with nail polish.

### Immunohistochemistry (IHC) staining

It is performed as described previously^37^. Briefly, fixed organoids are embedded in paraffin and sectioned. Paraffin sections are dewaxed and rehydrated by going through a ladder of xylene, ethanol and distilled water as described. Sections are then blocked and permeabilized with section staining buffer (5% FBS/0.2% gelatin/0.25% Triton X 100/PBS) for 30 minutes. Primary antibodies is diluted in section staining buffer at 1:400 and sections are stained at 4°C overnight. Sections are then washed with section staining buffer 3 times for 15 minutes each. Sections are then incubated with secondary antibodies diluted at 1:800 at 4°C overnight. After three round of washing, DAB substrate was eventually added to incubate the slides for 10 minutes. Slides are washed 3 times with PBS before mounting in 70% glycerol/PBS and sealed with nail polish.

### Quantification of anti-Cleaved Capase-3

We analyzed the image by Fiji software v2.14.0. We define the apoptosis index (%) as the anti-cleaved caspase-3 area divided by the organoid area. Unpaired student-T student is performed by Graphpad Prism 9.5.1. At least five regions randomly from three organoids were selected for quantification. ^**^ p<0.01.

### Flow cytometry

Vascular immune organoids (VIOs) were washed with PBS 3 times and were dissociated by 30 mins incubation of accumax (StemCell Technologies, 07921). Cells were washed once and resuspended in 100μl blocking buffer (2%FBS and 1% FcX (BioLegend, 422302) in PBS) at room temperature for 10 mins. Optimal concentration of CD144-PE (BD, 561714) and CD31-PE-Cy7 (BD, 563651) were added into the blocking buffer and were incubated for 30 mins on ice. Cells were washed with PBS once, followed by a centrifugation at 500g for 5 mins to remove supernatant.Cells were resuspended in sufficient staining buffer (2% FBS in PBS) with DAPI. Analysis were performed on a BD FACSymphony A3 cell analyzer. Single cells were gated based on FSC-A vs FSC-H, followed by the removal of dead cells by removing DAPI+ cells.

### SARS-CoV-2 infection

The infection was carried out in the designated facility in the University of Hong Kong following institutional safety measure (PMID: 36495872). In brief, VO, VIO, or isolated immune cells was exposed to a titter of 1E7 PFU/mL of SARS-CoV-2 virus. Samples were harvested 72 hours post-infection for either lysis for RNA collection or fixation for paraffin IHC. The SARS-CoV-2 used in this study is SARS-CoV-2 WT (HKU-001a, GenBank: MT230904), which was isolated from a clinical specimen of a COVID-19 patient.

### Single-cell RNA-seq

Single-cell RNA-seq was performed and analyzed as we published before^19^. Organoids were dissociated by accumax (STEMCELL Technologies, 07921). Briefly, We used accumax to dissociate organoids, and sorted out DAPI-negative cells by BD FACSAria™ Fusion. Library was prepared by Chromium Next GEM Single Cell 3’ Reagent Kit v3.1 as a service from the CPOS facility of HKU. A paired-end sequencing was performed by illumina NovaSeq 6000 and mapped to GRCh38 human reference genomics by CellRanger (v7.0.0). Seurat (v4.3.0) was used to filter out cells of lower than 500 features, 2000 counts, or above 15% mitochondrial content before calculating the read counts. Top 2000 highly variable genes were used to cluster cells by the uniform manifold approximation and projection (UMAP). Markers were identified by FindAllMarkers(), while cell types were annotated by well-known canonical markers.

### Bulk RNA-seq

Infected organoids or isolated immune cells were lysed for 10-30mins and collected RNA with Qiagen RNeasy Kits following the manufacturer’s protocol. Library was prepared by poly-A enrichment and sequenced by NovaSeq 6000 as a service provided by the CPOS of HKU. Raw reads were trimmed by Trim Galore and aligned to GRCh38.p14 human reference genome by STAR aligner. Aligned reads were converted into gene counts by samtools and featureCounts. Differentially expressed genes (DEGs) were analyzed by DESeq2 v1.30.1 in R v4.0.3. Gene set enrichment analysis (GSEA) was performed by gseGO() function from clusterProfiler v3.18.1. Volcano plots were generated by EnhancedVolcano v1.8.0. Representative gene expression was visualized by plotCounts() from DESeq2.

