## Supplementary figures for "SARS-CoV-2 infection activates inflammatory macrophages in vascular immune organoids"

#### Supplementary Figure 1.

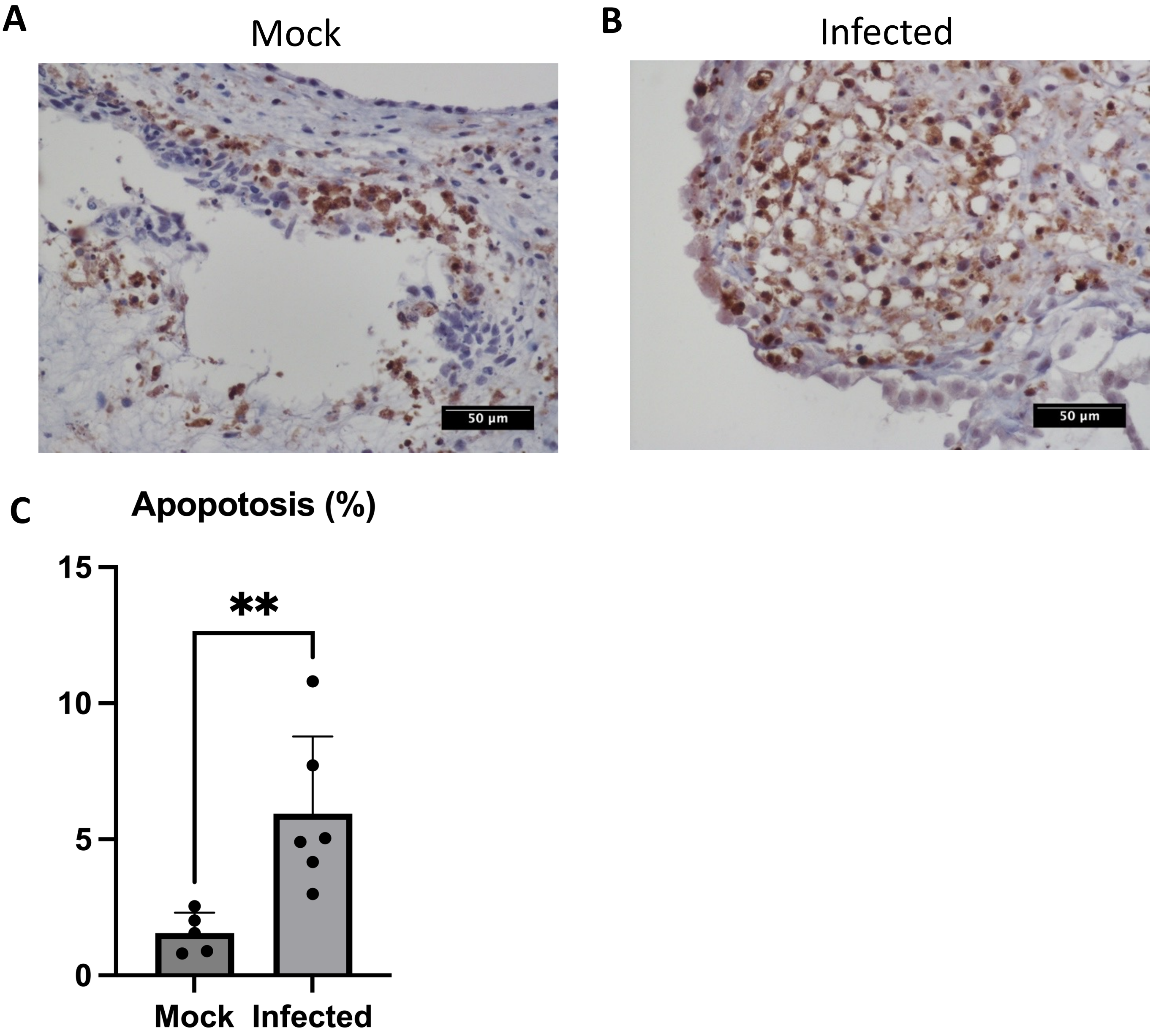


**Supplementary figure 1. Apoptosis of the infected VO.**

Immunohistochemistry staining of cleaved caspase-3 on **(A)** mock and **(B)** infected VOs. Scale bar = 50 µm. **(C)** Quantitative analysis of (A) and (B). ** $p < 0.01$.

**Supplementary Figure 2.**

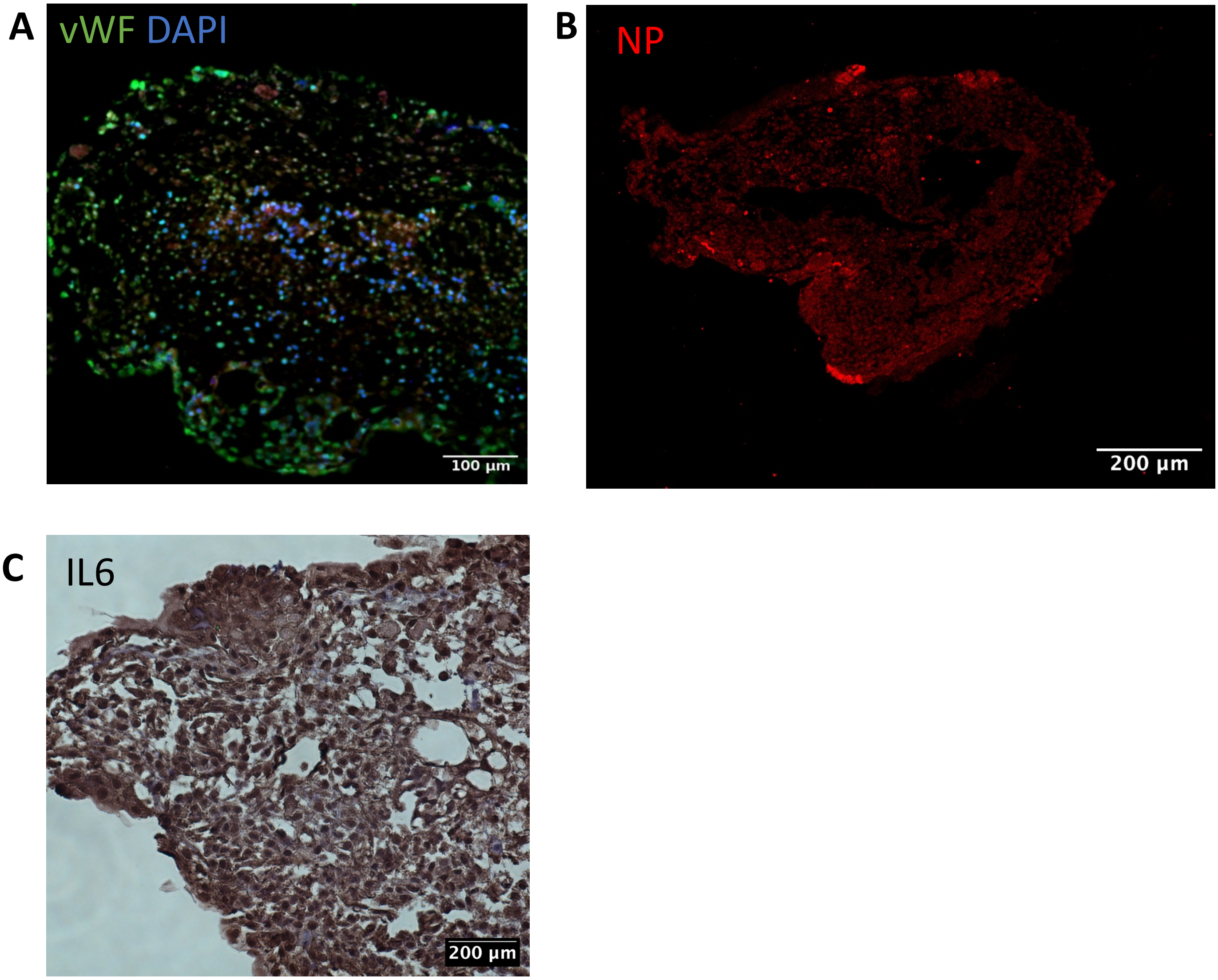


**Supplementary figure 2. Characterization of SARS-CoV-2 infected vascular organoids and vascular immune organoids. (A)** vWF immunofluorescent staining of VO. Scale bar = 100 µm. **(B)** N-protein (NP) immunofluorescent staining of VIOs. **(C)** IL6 immunohistochemistry staining of VIOs. Scale bar = 200 µm.

**Supplementary Figure 3.**

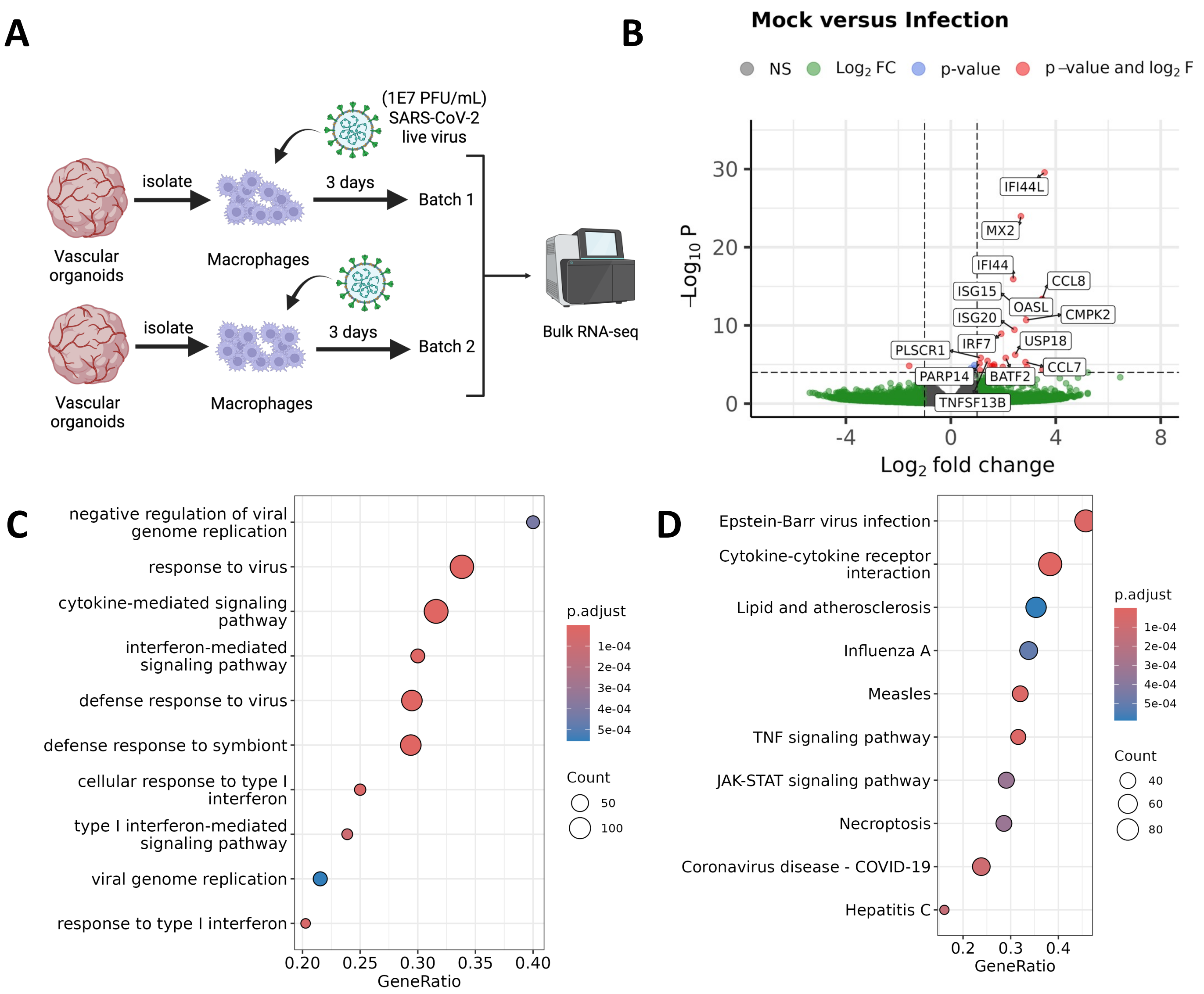


**Supplementary figure 3. SARS-CoV-2 activates inflammatory macrophages. (A)** Scheme of the SARS-CoV-2 infection on the isolated immune cells from the vascular immune organoids (VIOs). **(B)** Volcano plot , **(C)** Gene set enrichment analysis (GSEA), **(D)** and KEGG pathway analysis of the integrated VIO and isolated immune cells bulk RNAseq data.

### Supplementary Figure 4.

A

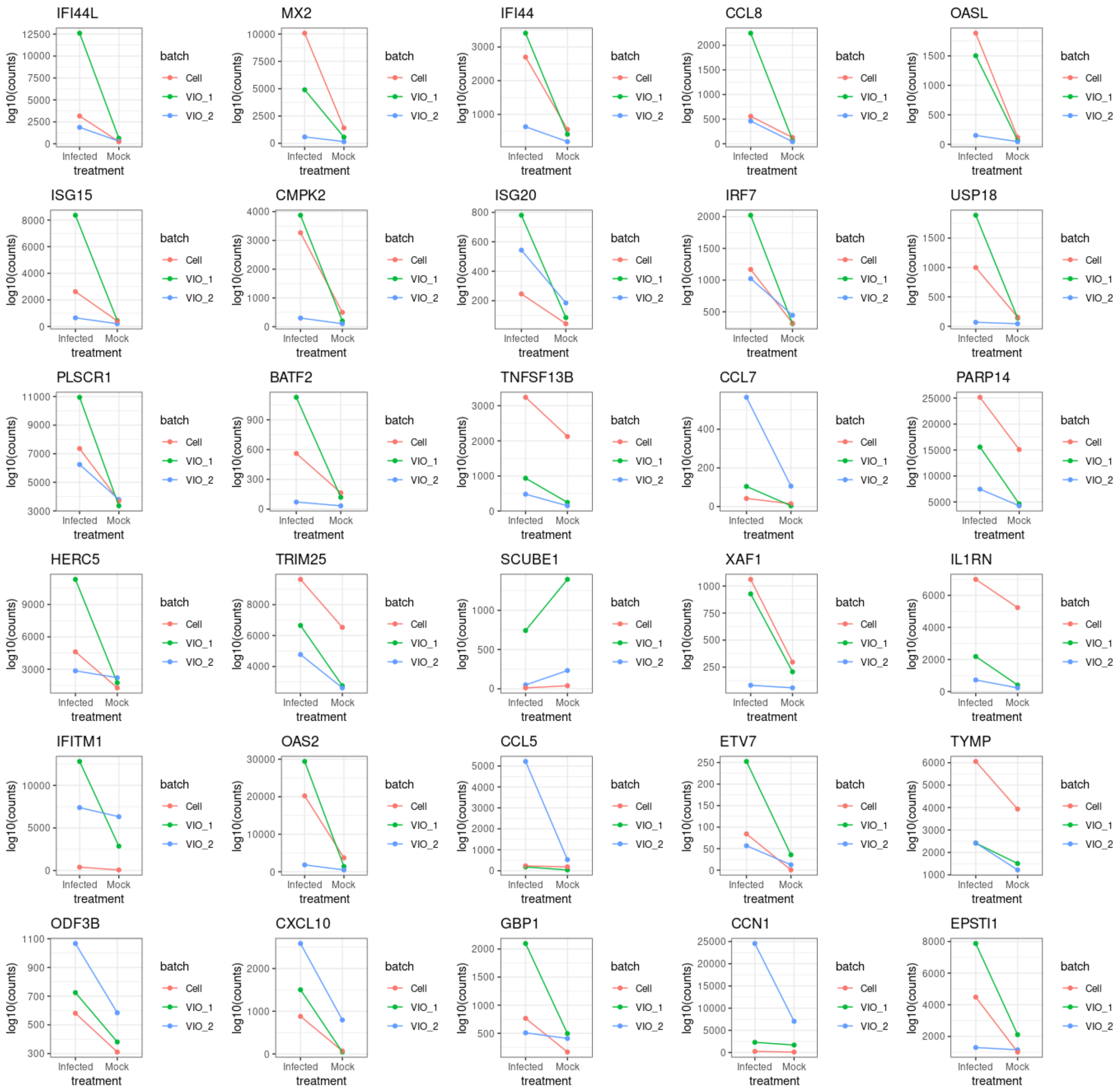


**Supplementary figure 4. Integrated analysis demonstrate mostly macrophage specific response to SARS-CoV-2 (A)** Batch-specific expression of top 30 differential expressed genes from the integrated analysis. “Cells” are the two batches of isolated macrophages. ”VIO_1” and “VIO_2” are two batches of VIO.
